# Copper acquisition in *Bacillus subtilis* involves Cu(II) exchange between YcnI and YcnJ

**DOI:** 10.1101/2025.05.23.655588

**Authors:** Yuri Rafael de Oliveira Silva, Grayson Barnes, Dia Zheng, Daniel Zhitnitsky, Samuel J. Geathers, Stephen C. Peters, Veronika A. Szalai, John D. Helmann, Oriana S. Fisher

## Abstract

The transition metal copper is biologically essential across all three domains of life. Several copper-dependent proteins and enzymes produced by the Gram-positive bacterium *Bacillus subtilis* have been characterized. However, many questions remain about how copper is recognized and trafficked to metalate cuproproteins. The *ycnKJI* operon in *B. subtilis* encodes a suite of proteins implicated in copper uptake and regulation, including the copper-binding protein YcnI and the putative copper importer YcnJ. Here, we demonstrate that one of the extracellular domains within YcnJ (YcnJ^CopC^) binds Cu(II) in 1:1 stoichiometry with high affinity using a histidine brace motif. Biochemical results reveal that YcnJ^CopC^ and YcnI can exchange Cu(II). Genetic studies reveal that loss of either YcnI or YcnJ, or mutation of the key residues required for Cu(II)-binding, leads to a growth defect under conditions of copper limitation. Together, these data suggest that the Cu(II)-binding sites in both YcnI and YcnJ contribute to efficient import under Cu limited conditions. Our results support a model in which YcnI may sequester Cu(II) from YcnJ, serving a regulatory role to limit the amount of copper that enters the cytoplasm and allowing Cu(II) to be stored for later import in the outer face of the membrane. This transfer of Cu(II) between extracellular domains of membrane-bound proteins represents a potential new paradigm in bacterial copper usage.

## Introduction

Bacteria require transition metals including iron, zinc, manganese, and copper. The *Bacillus subtilis* cytosol contains approximately 10^4^ free Mn and Fe ions, but virtually no free Cu (1–3). Most of the bacterial cytosolic Cu pools are bound to proteins (4, 5) and possibly low molecular weight thiols (6, 7). Although most Cu-containing enzymes function in the membrane or externally to the cell, most evidence suggests that protein metalation requires that Cu must first enter the cytoplasm before ultimately metalating membrane-bound or periplasmic cuproproteins (8). Under conditions of Cu sufficiency, some metal may enter the cell non-specifically, but when Cu is less abundant, dedicated high affinity Cu importers may be required (8). Surprisingly little is known about how extracellular copper is recognized and acquired by any bacterial species, including well-characterized model organisms such as the Gram positive bacterium *B. subtilis*.

In *B. subtilis*, genetic and physiological studies have implicated the *ycnKJI* operon in copper homeostasis. The *ycnKJI* operon is upregulated under copper starvation conditions and downregulated under copper replete conditions. A *ycnJ* deletion strain was reported to have a growth defect and reduced intracellular Cu accumulation under conditions of Cu limitation (9). These data suggested that the YcnJ protein may serve as a Cu importer. Proteins homologous to YcnJ have been implicated in similar roles; for example, the *copCD* genes in *Pseudomonas syringae* confer sensitivity to Cu and their presence results in increased intracellular Cu accumulation (10), and the YebZ protein from *E. coli* has also recently been implicated in copper uptake (11). Following import, Cu interacts with metalloregulatory proteins (CsoR, YcnK) that monitor Cu status (9, 12, 13) and is trafficked to support Cu-dependent metalloenzymes. During growth, the only known Cu-dependent enzymes are the major quinol oxidase (Qox complex) and cytochrome *c* oxidase (Cta complex) in the cell membrane. During sporulation, a Cu-dependent laccase (CotA) is synthesized that is part of the spore coat (14).

The YcnJ protein has two extracellular domains (an N-terminal CopC domain and a C-terminal YtkA domain) that sandwich a membrane-embedded CopD domain. Proteins containing CopD domains are widespread throughout bacterial phylogeny and those in several organisms, including YcnJ in *B. subtilis* (9, 11, 13), have been proposed to serve as Cu importers based on microbiological data and amino acid sequence similarity (15, 16). The domain architecture and genome neighborhood for YcnJ from *B. subtilis* differs from the most common *copD* sequences found across bacterial phylogeny (11), as YcnJ notably is encoded by a gene located between *ycnI* and *ycnK* and is a CopC-CopD-YtkA fusion. In addition to the YtkA domain in YcnJ, *B. subtilis* has another YtkA paralog that functions in metal-loading of a cytochrome *c* oxidase (17). No other YtkA domains have been investigated experimentally, although homologs of YtkA are found in many *Bacillus* species as well as within other bacterial species, predominantly within *Pseudomonadota* (18). CopC proteins are found primarily in bacteria and are classified according to the presence of Cu(I) and Cu(II) binding sites. Alignment of the amino acid sequence of the N-terminal CopC domain of YcnJ to representative sequences of each subfamily of CopC proteins suggests that this domain falls into a class (denoted C_0-1_) that contains a single Cu(II) binding site coordinated by histidine and aspartic acid residues (**Fig. 1b**) (16). In some organisms CopC domains exist as single-domain proteins, but in other cases, as in *B. subtilis*, they are fused to other domains (16, 19–21). The biological role of YcnJ^CopC^ has not been determined.

**Figure 1.**
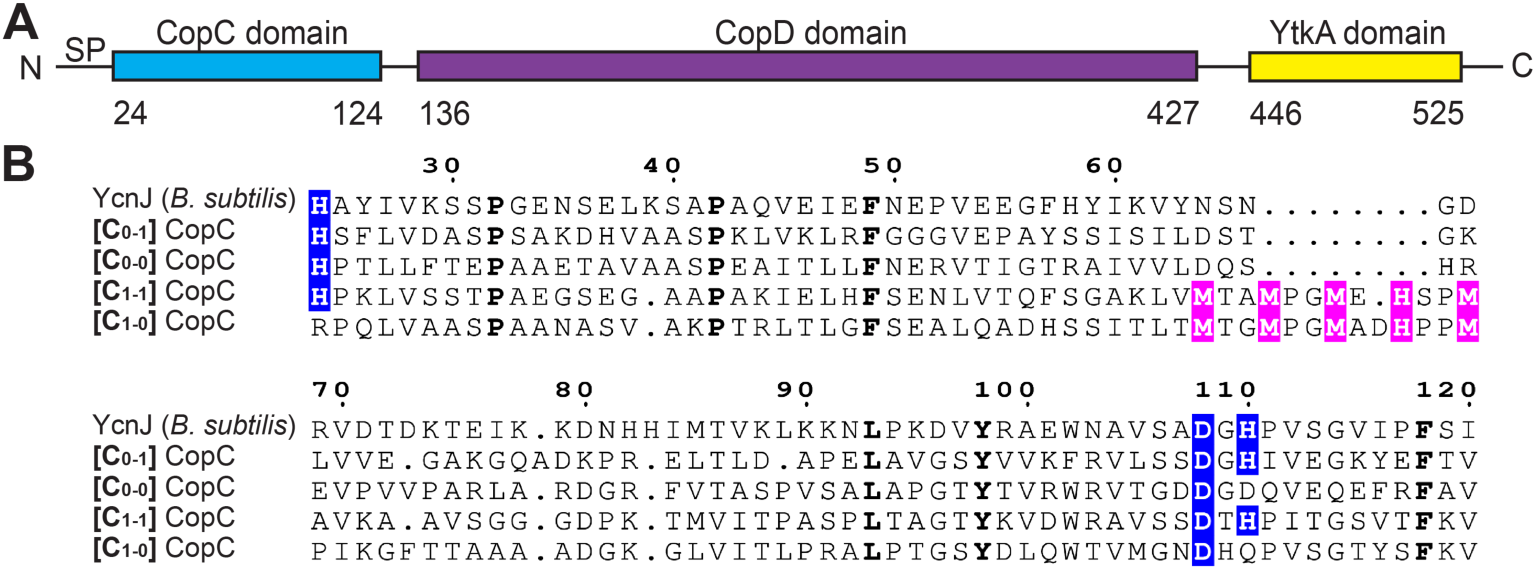
The CopC domain of YcnJ is a [C_0-1_]-subfamily CopC. (a) Domain architecture of YcnJ from *B. subtilis*. (b) Sequence alignment between the CopC domain of YcnJ and representatives from each of CopC subfamilies. Residues involved in Cu(II) coordination are highlighted in blue, while those involved in Cu(I) coordination are highlighted in pink. Residues with 100 % conservation are presented in bold. [C_0-1_] CopC: *Methylosinus trichosporium* OB3b (WP_003610846.1), [C_0-0_] CopC: *Mycolicibacterium tusciae* (WP_239591552.1), [C_1-1_] CopC: *Pseudomonas syringae* pv. tomato (WP_003317555.1), [C_1-0_] CopC: *Sphingobium yanoikuyae* XLDN2-5 (WP_010339668.1).

The *ycnKJI* operon also encodes the functionally uncharacterized YcnI protein that, like YcnJ, is localized to the membrane. YcnI contains an extracellular domain of unknown function (DUF1775) that we previously discovered adopts a cupredoxin-like fold that binds Cu(II) with femtomolar affinity in 1:1 stoichiometry (22, 23). The YcnI Cu-binding site uses a monohistidine brace motif featuring an N-terminal histidine that uses both the amino terminus of the polypeptide chain and the imidazolium side chain to coordinate the Cu ion, a glutamate residue that coordinates the metal through its carboxylate side chain, and a highly conserved tryptophan residue (22). The latter is critical for proper and stable orientation of the metal ion within the binding site (23). Beyond its ability to tightly and specifically bind Cu(II), the function of YcnI has not been determined.

Just as in the *B. subtilis ycnKJI* operon, across bacterial phylogeny DUF1775 and CopC domains are frequently encoded by genes that neighbor genes encoding CopD domains (16, 22). This suggests that the DUF1775 and CopC domains, which are nearly always predicted to be periplasmic in Gram negative bacteria or fused to transmembrane domains in Gram positive organisms, may work in concert with the CopD domains to facilitate or regulate Cu uptake across the membrane.

To investigate the role of YcnJ and YcnI in *B. subtilis*, here we structurally and spectroscopically characterize YcnJ^CopC^. We find that it binds Cu(II) through a histidine brace motif with sub-femtomolar affinity, similar to the binding mode we reported for the DUF1775 domain of YcnI (22). The similarities in the coordination environment, stoichiometry, binding affinity, and Cu oxidation state preferences led us to investigate potential interplay between these two extracellular domains. We find that Cu(II) can be transferred between these domains *in vitro* and that *in vivo* point mutations at the metal-binding sites in YcnI and YcnJ *B. subtilis* phenocopy *ycnI* and *ycnJ* deletion strains. Together, these data provide evidence that extracellular interactions with Cu(II) contribute to growth of *B. subtilis* under Cu-limiting conditions. These results further our understanding of how Cu is recognized and acquired in *B. subtilis* and suggest that other bacterial species may also use homologous proteins in analogous ways.

## Results

### Structure of the extracellular CopC domain of YcnJ

YcnJ from *B. subtilis* has been implicated as a Cu importer, proposed to use its membrane-bound CopD domain for Cu uptake (9, 13). The full-length protein, however, includes additional extracellular domains: a CopC domain at its N-terminus and a YtkA domain at its C-terminus, whose roles in this process have not been investigated (**Fig. 1a**). All characterized members of the CopC family can either bind a single Cu(II) at an N-terminal site, a single Cu(I) using a methionine-rich region, or both; sequence analysis of family members suggests that nearly all CopC domains can be classified according to the presence and coordination environment of these Cu-binding sites (16, 21). These classes are denoted by a pair of subscript numerals, where the first number represents the absence (0) or presence (1) of the Cu(I) site ligands and the second number represents the absence (0) or presence (1) of a Cu(II) site coordinated by His ligands (**Fig 1b**) (16, 21). Alignment of the amino acid sequence of the N-terminal CopC domain of YcnJ to representative sequences of each subfamily of CopC proteins suggests that this domain falls into the C_0-1_ class due to the conservation of the two histidines and one aspartate used to form the Cu(II) binding site and the absence of a predicted Cu(I)-binding motif (**Fig. 1b**).

We initiated structural studies to investigate how this domain compares to other characterized CopC domains. The full-length YcnJ includes a predicted signal peptide (24); assuming that this portion of the sequence is cleaved post-translationally, there is a His residue at the native N-terminus of the protein. We designed a construct of the CopC domain with an N-terminal His-SUMO tag, allowing us to generate the native N-terminus by cleavage with Ulp1 protease. We purified the resulting YcnJ^CopC^ protein and loaded it with Cu(II) to try to obtain crystals in its Cu-loaded form. However, we were only able to successfully crystallize the as-purified sample. We determined its structure to 0.160 nm (1.60 Å) resolution (**Fig. 2a, Table S1**). The overall fold displays the typical β-barrel structure of a cupredoxin fold observed in other CopC proteins (16, 25). YcnJ^CopC^ has a root mean square deviation (RMSD) of 0.136 nm to 0.183 nm (1.36 Å to 1.83 Å) between its structure and those of other CopC homologs. Of those CopC homologs whose structures have been experimentally determined, YcnJ^CopC^ has the lowest RMSD with the Cu(II)-binding CopC protein from *M. trichosporium* OB3b (**Fig. 2b, Fig. S1**) (16, 21, 26). The main difference between YcnJ^CopC^ and these other proteins is a disruption of β4. This could, however, be a crystallization artifact since we also observe electron density for triethylene glycol and a sulfate ion from the crystallization solution near this beta strand (**Fig. S2**).

**Figure 2.**
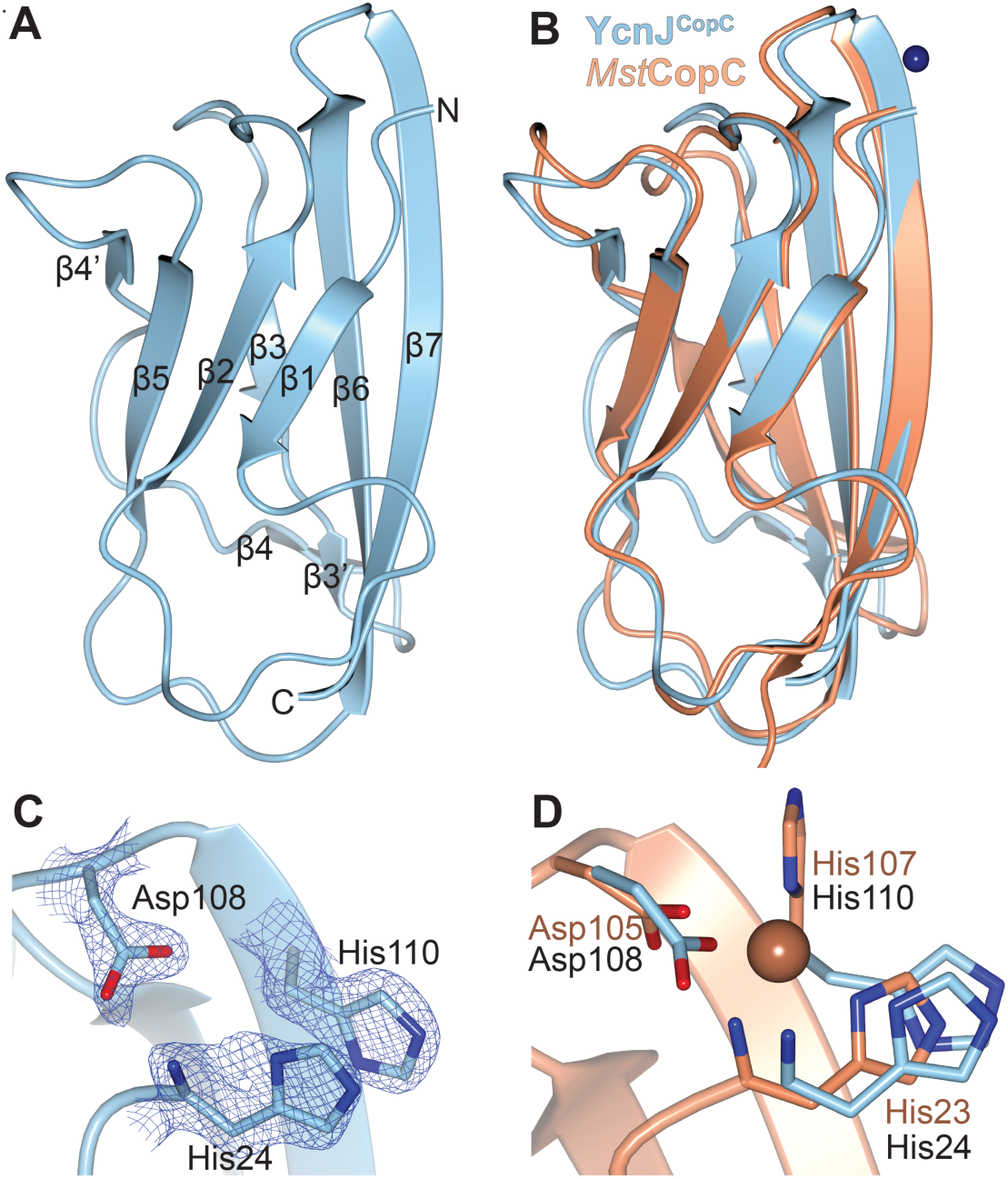
Structure of YcnJ^CopC^. (a) Crystal structure of YcnJ^CopC^ and (c) close-up view of its putative Cu(II) binding site with 2*F_o_-F_c_* map in blue mesh contoured to 1.5 σ. (b) Superposition of YcnJ^CopC^, in blue to Cu(II)-bound *Methlyosinus trichosporium* CopC, in orange (PDB ID 5ICU (16)) and (d) close-up view of its Cu(II)-binding site. The Cu(II) ion is shown as a sphere.

In CopC proteins that bind a single Cu(II) ion, an N-terminal His residue coordinates the metal by both the amino terminus and the δ-nitrogen from the side chain (**Fig. S1**) (16). Another His and an Asp from a DxH motif provide a third coordinating nitrogen and a coordinating oxygen, respectively. These three residues are conserved in YcnJ^CopC,^ and the crystal structure shows that these amino acids are close to one another, suggesting that they may be involved in Cu-binding similar to other metalated proteins in this family (**Fig. 2c, d; Fig. S1**). Despite the similarities to other experimentally characterized CopC proteins, which to date have all been single-domain proteins, YcnJ is different because it is a multidomain protein, raising additional questions about how its CopC domain may work in concert with its other domains.

### YcnJ^CopC^ binds Cu(II) in a 1:1 stoichiometry

To determine how YcnJ^CopC^ might bind Cu, we turned to spectroscopic approaches. First, we loaded as-purified protein samples with 1 or 2 molar equivalents of Cu(II), and measured copper concentrations by inductively-coupled plasma-mass spectrometry (ICP-MS). The resulting data show approximate 1:1 stoichiometry for the Cu:YcnJ^CopC^ complex (**Fig. 3a**). To confirm the oxidation state of the metal, we analyzed Cu-loaded YcnJ^CopC^ by electron paramagnetic resonance (EPR) spectroscopy. We observe a spectrum characteristic of axial Cu(II) coordination in the Cu-loaded sample, with parameters of g_||_ = 2.257, g_⊥_ = 2.055, and A = 17.86 mT. On the other hand, we do not observe spectral features in our as-purified protein, indicating that this sample does not contain any paramagnetic species (**Fig. 3b**). We also observe seven distinct hyperfine lines, which could arise from the presence of three geometrically equivalent nitrogens; this is consistent with each of the three nitrogens from the amino terminus, His24, and His110 side chains coordinating the metal ion (**Fig. 3c**).

**Fig. 3.**
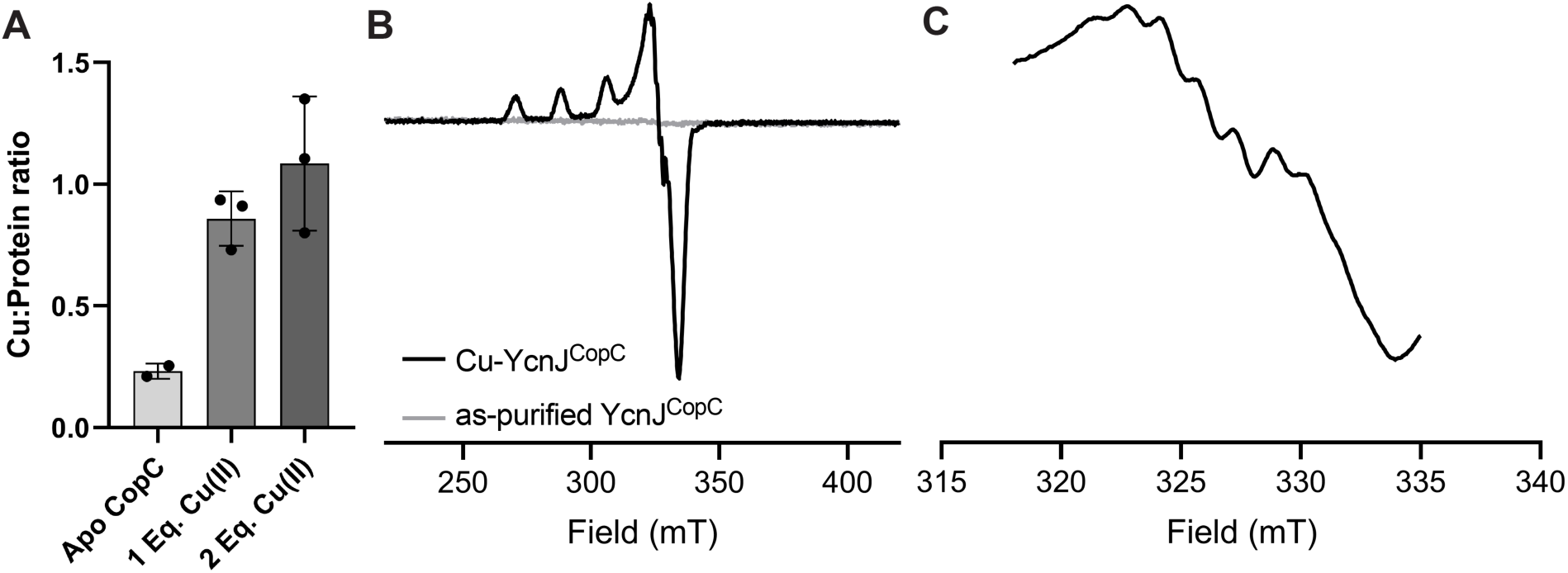
YcnJ^CopC^ binds Cu(II). (a) Cu-binding stoichiometry of the CopC domain determined by ICP-MS, indicating it binds only Cu(II) in a 1:1 stoichiometry. n = 2 for as-purified, n = 3 for Cu-loaded samples. Data represent mean ± SD of biological replicates. (b) CW EPR spectra of Cu-YcnJ^CopC^ (black) and as-purified (grey) collected at 80 K with a microwave power of 0.4 mW, 4 scans. (c) Narrow spectrum showing the superhyperfine lines for Cu-YcnJ^CopC^ on the g_⊥_ peak of the spectrum shown in (b). Microwave power is 0.4 mW, 4 scans. Seven superhyperfine lines are clearly discernible.

The CopC domain of YcnJ is strikingly similar to the extracellular DUF1775 domain of YcnI (YcnI^DUF1775^), the protein of unknown function encoded by the *ycnI* gene that is immediately downstream of *ycnJ* (**Fig. 6b, c**). YcnI^DUF1775^, like YcnJ^CopC^, is a small globular domain that binds a single Cu(II) ion with femtomolar affinity (22, 23), coordinating it through two nitrogens from its N-terminal histidine and an oxygen provided by a negatively charged residue. YcnI lacks the second coordinating histidine, instead stabilizing the Cu(II) binding site through interactions with a tryptophan residue (19). We therefore decided to measure the binding affinity of YcnJ^CopC^ for Cu. We conducted ITC using glycine, a weak Cu(II) chelator, as a competitor for copper to allow us to obtain more accurate affinity values in the measurable range for such a tight interaction (50) (**Fig. 4, Fig. S3**). The conditional *K*_D_ calculated was 1.43 × 10^-16^ mol/L, an order of magnitude tighter than the affinity of YcnI for Cu(II) (23), but similar to the metal affinities that have been reported for other proteins in the CopC family (20, 26) (**Table S3**). The thermodynamic properties of the association of YcnJ^CopC^ for Cu(II) are also enthalpically favorable (ΔH = −77 487 J/mol (−18 520 cal/mol); ΔS = −109.6 J/mol/K (−26.2 cal/mol/K)), similar to what we observed for YcnI under identical conditions with the same glycine concentration as a weak competitor (23).

**Figure 4.**
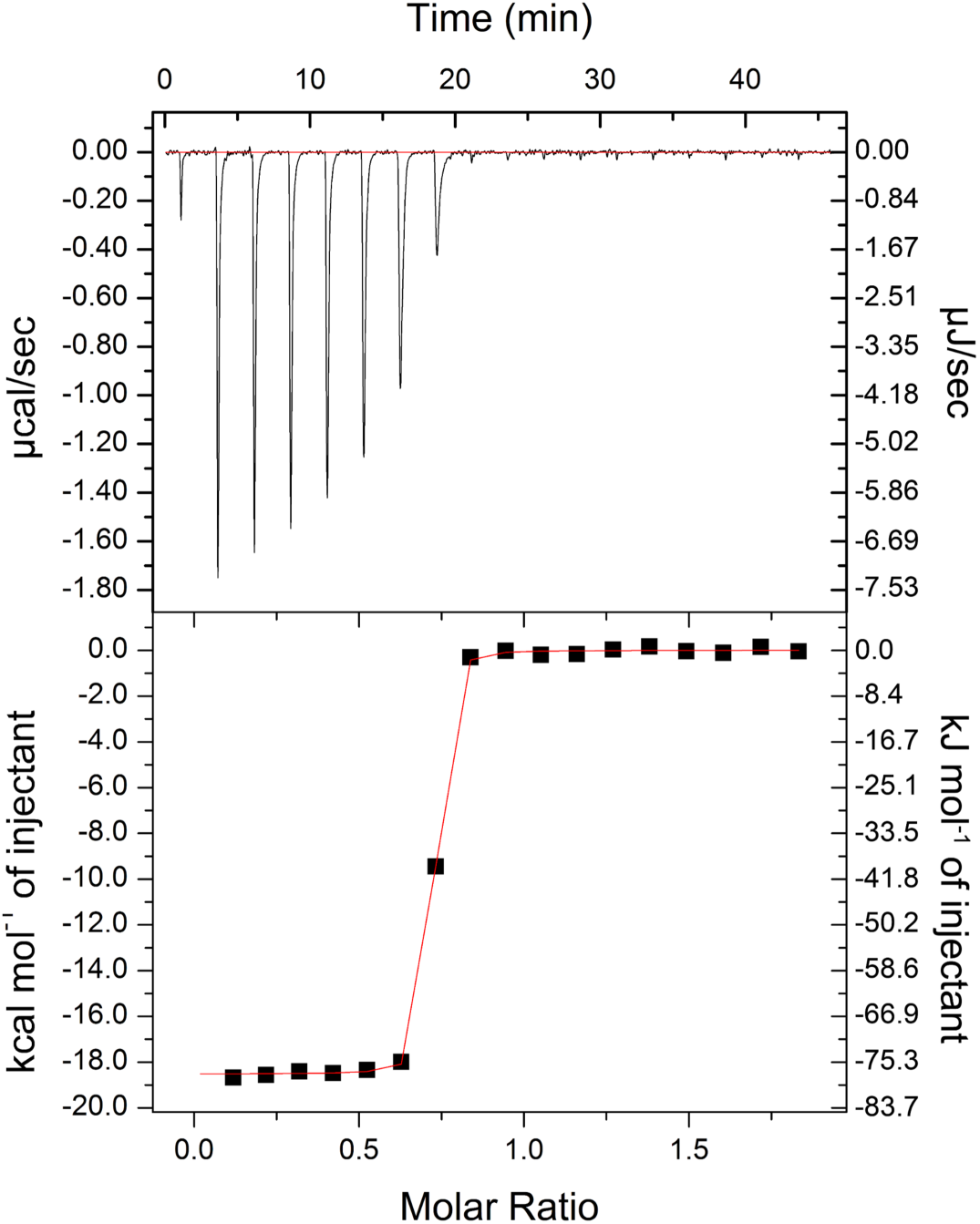
Isothermal titration calorimetry of Cu(II) into YcnJ^CopC^. Measurements were performed in the presence of 30 mmol/L glycine as a weak competitor. Representative thermogram (Refer to **Fig. S3** for replicate). Conditional *K*_D_ = 1.43 (± 0.06) × 10^-16^ mol/L; n_ITC_ = 0.682 ± 0.0007; ΔH = −77 487 ± 216 J/mol; ΔS = −109.6 J/mol/K. Reported error represents error on the curve fit.

The stoichiometry (n_ITC_) for the Cu(II) interaction with YcnJ^CopC^ is less than one. Similarly sub-stoichiometric values for n_ITC_ have been reported for other high-affinity Cu-binding proteins, including other CopC homologs (27–30). One possible explanation for nITC < 1 is that the high affinity could mean that the protein becomes partially metalated during purification or that there is a subpopulation of YcnJ^CopC^ that cannot bind metal at all.

### In vivo role of YcnI and YcnJ

Considering our *in vitro* findings that both YcnI and YcnJ contain extracellular Cu(II) binding sites, we decided to investigate the effects of Cu(II) levels physiologically. We generated markerless gene deletion *B. subtilis* strains for both genes (*ΔycnI* and *ΔycnJ*) and compared growth of these strains to the wild-type organism in minimal media with or without the Cu(II)-specific chelator triethylenetetramine (TETA). Similar to previous studies (9), we do not observe significant differences among cells grown in minimal medium with malate as the carbon source in the absence of an added Cu chelator (**Fig. 5A**). However, in the presence of 10 mmol/L TETA we observe that growth of the parent strain (WT) slows and the highest cell yield (as judged by optical density at 600 nm; OD_600_) that is achieved is reduced related to the untreated cultures. Upon cessation of growth, the OD_600_ decreases slowly over time, as is commonly observed with *B. subtilis*. This decrease is due to autolysis of cells and is exacerbated when cells are depleted of energy and there is a decrease in proton motive force (31). Compared to the parent strain, the *ΔycnJ* mutant strain has a reduced growth rate and >2-fold reduced cell yield in the presence of TETA (**Fig. 5B**). In contrast, the Δ*ycnI* strain had a similar growth rate as the parent strain in the presence of TETA, but the final cell yield was reduced (**Fig. 5C**). While a growth defect was anticipated for the *ΔycnJ* strain (9), a phenotype for *ycnI* mutants had not previously been reported.

**Figure 5.**
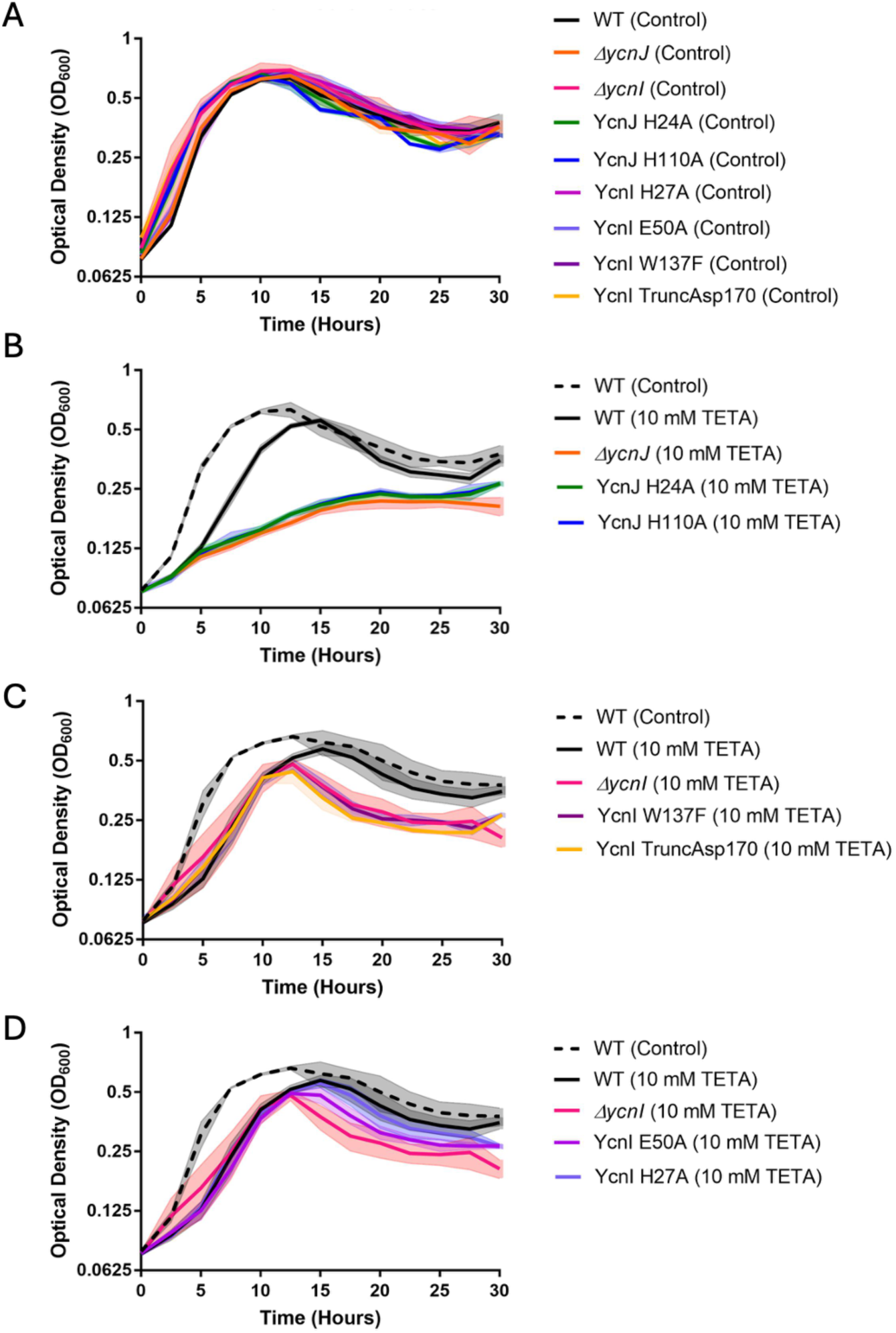
Growth curves for *B. subtilis* strains. Growth curves of various *B. subtilis* strains in (A) minimal malate medium with no chelator. (B) With 10 mmol/L TETA the Δ*ycnJ* mutant and the *ycnJ* His24Ala and His110Ala alleles are severely growth compromised. (C) With 10 mmol/L TETA the Δ*ycnI*, *ycnI* Trp137Phe and *ycnI* Asp170* (transmembrane helix deletion) alleles all have a comparable decrease in growth yield. (D) With 10 mmol/L TETA the *ycnI* His27Ala and *ycnI* Glu50Ala alleles have a reduced growth defect compared to the Δ*ycnI* null mutant. Note that some curves are repeated between panels for ease in comparison. Shaded areas report the standard deviation of 3 biological replicates.

Given that both *ycnI* and *ycnJ* appear to support growth under Cu(II) limitation, we next hypothesized that this phenotype may be directly linked to the extracellular Cu(II)-binding sites in YcnJ^CopC^ and YcnI^DUF1775^. We generated *B. subtilis* strains in which we introduced single point mutations at the Cu(II) binding site in the YcnI^DUF1775^ monohistidine brace motif (Trp137Phe, His27Ala, Glu50Ala), in the YcnJ^CopC^ histidine brace (His24Ala, His110Ala), and also removed the anchoring C-terminal transmembrane helix of YcnI (Asp170*). Strikingly, we find that mutation of either histidine in YcnJ^CopC^ phenocopies the *ΔycnJ* strain when grown in TETA-treated media (**Fig. 5B**), suggesting that Cu(II)-binding to the CopC domain is required for YcnJ function. Similarly, mutation of either the Trp137Phe YcnI protein, and YcnI lacking its C-terminal transmembrane helix have nearly identical growth curves to the *ΔycnI* strain in media treated with TETA (**Fig. 5C**), suggesting that YcnI requires both the Cu-binding site as well as membrane localization. Similarly, the Glu50Ala YcnI protein also had a reduced growth yield (**Fig. 5f**). In contrast, the His27Ala mutation in YcnI had a relatively small effect (**Fig. 5D**). This mutation removes the coordinating nitrogen from the histidine side chain but retains the nitrogen ligand from the amino terminus of the peptide backbone.

### Cu(II) can be transferred between YcnI^DUF1775^ and YcnJ^CopC^ in vitro

The finding that both YcnJ and YcnI help support growth under Cu limitation (and that this role requires residues that bind Cu) together with previous studies that indicated that YcnI and YcnJ are under the control of a shared promoter and are both membrane-localized (9, 13) led us to hypothesize that YcnI might deliver Cu to YcnJ through its CopC domain. The coordination environment of the Cu(II) bound to YcnJ^CopC^ and that of YcnI^DUF1775^ both involve an N-terminal His that provides two ligands to the metal ion in addition to a negatively charged side chain from either an Asp or a Glu (22, 23). Notably, however, the Cu(II) in YcnI^DUF1775^ has fewer protein-derived ligands than the Cu(II) in typical CopC structures (16, 21). In most CopC proteins, the Cu ion is in a relatively solvent-exposed region of the domain (**Fig. 6a,b**). On the other hand, in its apo form, YcnJ^CopC^ has π-π stacking between the two imidazole rings of the two conserved His side chains, an interaction not accessible to YcnI because it lacks a second His ligand (**Fig. 6c,d**). Additionally, in both proteins, the Cu-binding sites are solvent accessible.

**Fig. 6.**
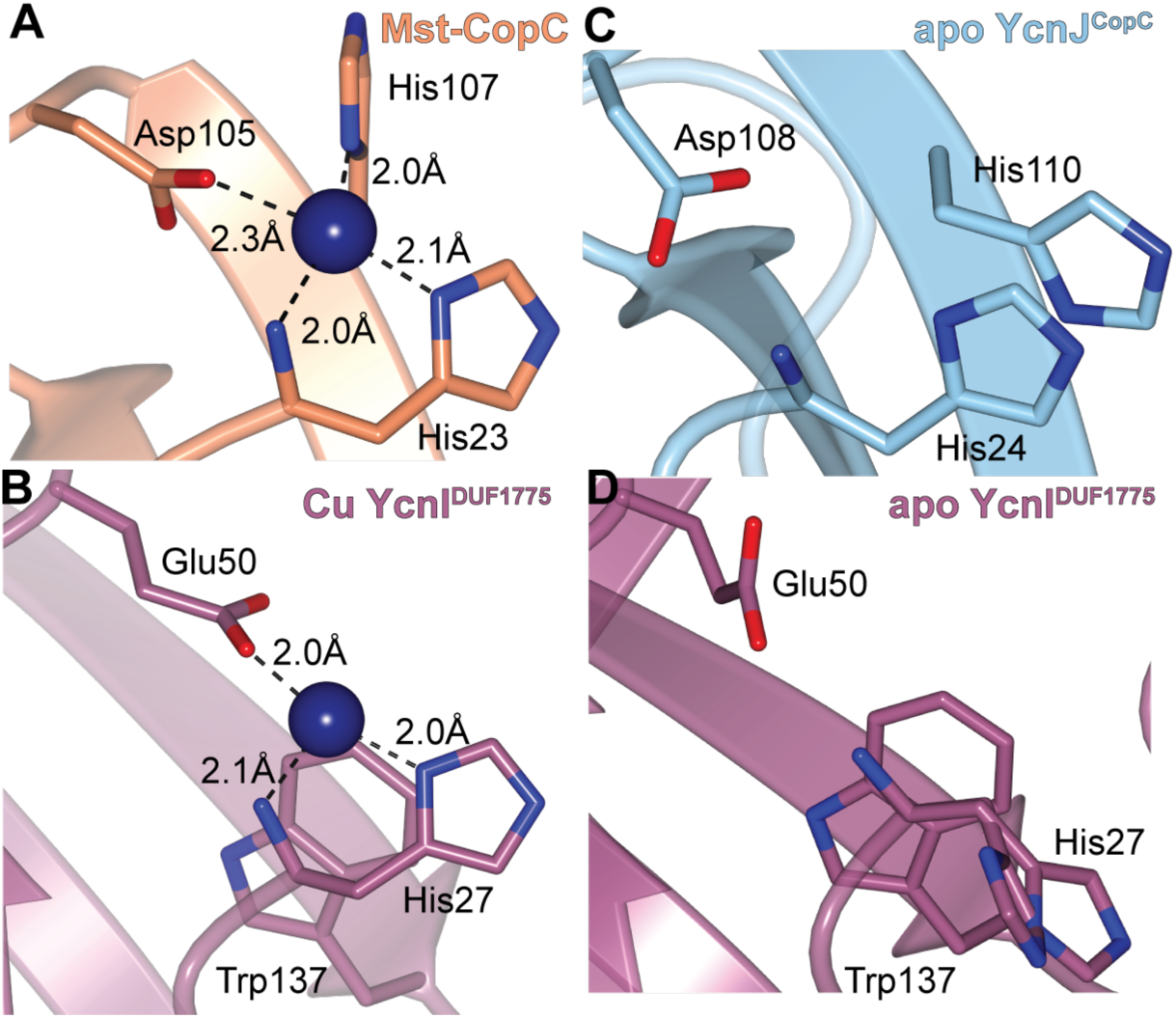
Structural comparison of Cu-bound and apo YcnJ^CopC^ and YcnI^DUF1775^. (a) Cu(II)-binding site of *Methylosinus trichosporium* OB3b CopC (PDB ID: 5ICU) and (b) Cu(II)-binding site in YcnI^DUF1775^ (PDB ID: 7MEK) (22) compared to the apo structures of (c) YcnJ^CopC^ (PDB ID: 8UM6, this study) and (d) YcnI^DUF1775^ (PDB ID: 7ME6) (22).

YcnI and YcnJ are encoded by the same operon, localized on the extracellular side of the membrane, and both have solvent accessible Cu-binding sites. We therefore considered the possibility that perhaps they could act as metallochaperones to shuttle Cu ions between them, analogous to other bacterial Cu-binding proteins that facilitate transfer of copper ions to their partner proteins (32–37). Copper transfer between proteins often occurs *via* weak, transient interactions between binding partners, at times with affinities weaker than those that can readily be measured by traditional protein-protein interaction assays (7, 38–41).

To monitor transfer from YcnJ to YcnI, we incubated Cu-loaded YcnJ^CopC^ with apo YcnI^DUF1775^, chromatographically separated them, and measured the Cu content of the resulting proteins (**Fig. 7a**). Across multiple independent experiments, we consistently observe a decrease in the Cu content of WT YcnJ^CopC^ and a corresponding increase in Cu content of WT YcnI^DUF1775^. To confirm that this effect is not due to loss of Cu from YcnJ^CopC^ into buffer during the transfer experiment, we also conducted a control experiment by incubating Cu-loaded YcnJ^CopC^ in buffer; although we observe a small decrease in Cu content after chromatography, it is more modest than in the transfer experiments (**Fig. 7c**).

**Figure 7.**
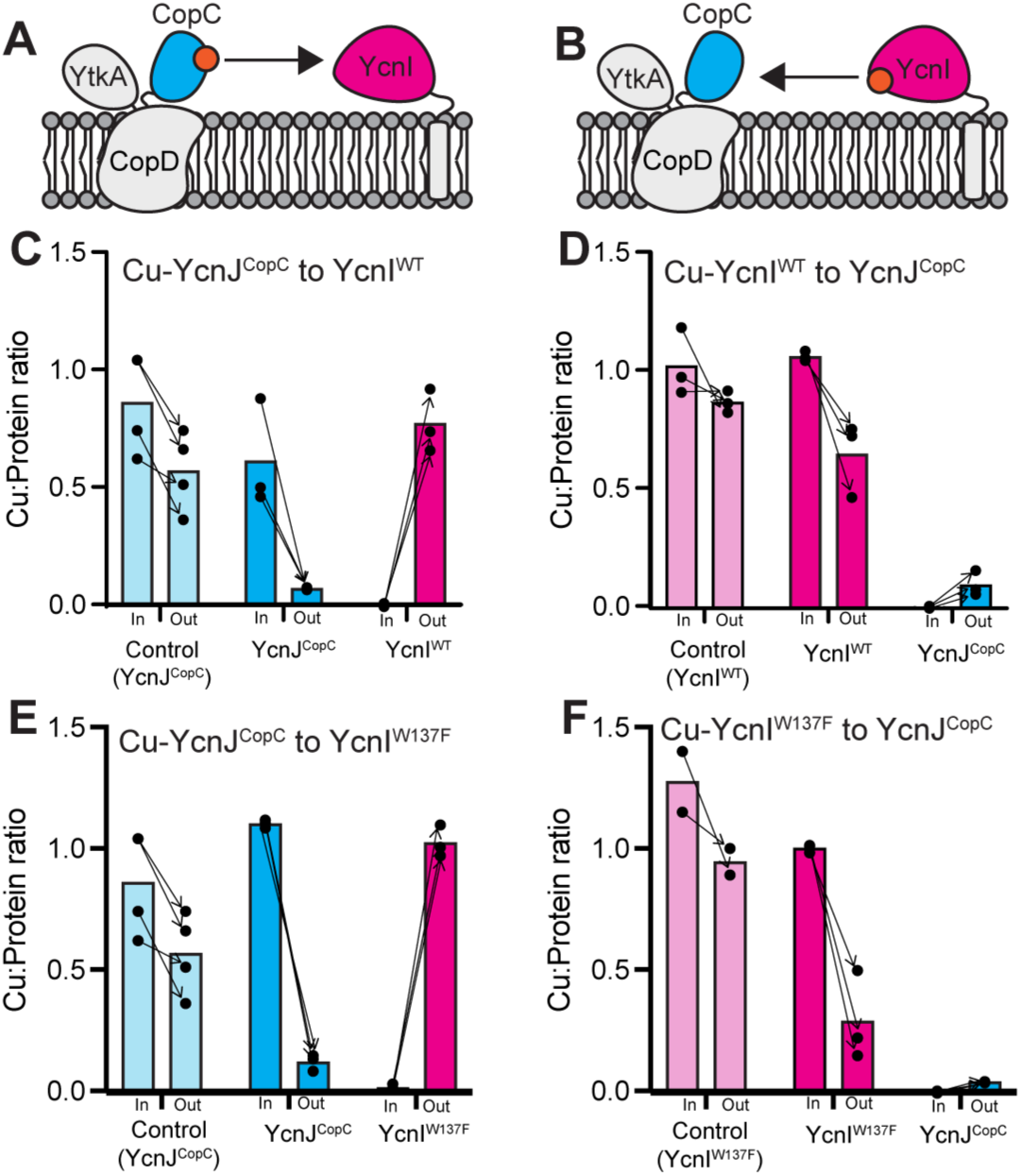
Copper transfer between YcnI and YcnJ^CopC^. Cartoons depicting the two proposed mechanisms for Cu(II) transfer: (a) from YcnJ^CopC^ to YcnI and (b) from YcnI to YcnJ^CopC^. (c-f) Cu:protein ratios before and after Cu-transfer experiments. (c) Cu-YcnJ^CopC^ (blue) to apo WT YcnI^DUF1775^ (pink); light blue bars represent a Cu-YcnJ^CopC^ control in buffer, without a transfer partner. (d) Cu-WT YcnI^DUF1775^ (pink) to apo YcnJ^CopC^ (blue); light pink bars report average Cu:protein ratio of a Cu-WT YcnI^DUF1775^ control in buffer, without a transfer partner. (e) Cu-YcnJ^CopC^ to apo Trp137Phe YcnI^DUF1775^, colored as in part (c). (f) Cu-Trp137Phe YcnI^DUF1775^ to apo YcnJ^CopC^, colored as in part (d). Individual data points for each biological replicate are shown as black circles, with arrows indicating input and output from each individual experiment. Bars represent the average Cu:protein ratio across all replicates.

This indicates that, *in vitro*, Cu(II) can be transferred directly from YcnJ^CopC^ to WT YcnI^DUF1775^. To investigate whether Cu can be transferred from YcnI to YcnJ, we conducted the reverse experiment by incubating Cu-loaded WT YcnI^DUF1775^ with apo YcnJ^CopC^ (**Fig. 7b**). We do not observe a consistent reduction in the Cu content of WT YcnI^DUF1775^ in the control experiment, but we do see a consistent decrease in the Cu content of WT YcnI^DUF1775^ and a corresponding increase in that of YcnJ^CopC^ across multiple independent experiments (**Fig. 7d**). To address the possibility of passive Cu(II) transfer between YcnJ^CopC^ and YcnI, *i.e.* without direct interaction between the proteins, Cu-transfer experiments were repeated with proteins separated by a dialysis membrane. No significant copper loss in Cu(II)-loaded proteins was detected and no detectable Cu(II) binding was observed by the as-purified proteins (**Fig. S4**), supporting a requirement for physical interactions between YcnJ^CopC^ and YcnI, to allow Cu(II) transfer. Overall, we observe protein-mediated Cu(II) transfer in both directions, although perhaps more significantly from YcnJ^CopC^ to YcnI, suggesting that Cu(II) may be transferred preferentially from YcnJ to YcnI.

We also performed analogous experiments with a Trp137Phe YcnI mutant (Trp137Phe YcnI^DUF1775^) that we had previously found to have weaker affinity to Cu(II) than the WT protein and a more labile Cu(II)-binding site (23). Similar to our data with the WT YcnI^DUF1775^, we observe transfer of Cu(II) from YcnJ^CopC^ to Trp137Phe YcnI^DUF1775^ (**Fig. 7e**) and from Trp137Phe YcnI^DUF1775^ to YcnJ^CopC^ (**Fig. 7f)**. When YcnI is used as a donor, there is a decrease in its Cu:protein ratio post-transfer. This decrease is more noticeable in the Trp137Phe variant than the WT protein, potentially due to the decreased affinity of this mutant for Cu(II) (23). This suggests that Trp137 may not be essential for the transfer process but perhaps plays a different or more nuanced role in modulating Cu(II) recognition and acquisition in *B. subtilis*.

## Discussion

Our results suggest a role for extracellular Cu(II) binding in copper homeostasis in *B. subtilis*. We propose that the initials steps in copper acquisition occur when YcnI^DUF1775^ or YcnJ^CopC^ bind Cu(II). We find that the extracellular CopC domain of YcnJ binds Cu(II) tightly and that this interaction is important for growth under Cu-limiting conditions. The phenotypes of strains with single point mutations at the metal-binding sites in *ycnJ* and *ycnI* are generally similar to those of null mutants. The parallels in the Cu(II) coordination sites in YcnI^DUF1775^ and YcnJ^CopC^ led us to hypothesize that these domains may facilitate Cu(II)-transfer. Our *in vitro* experiments support this hypothesis and suggest that each domain can transfer Cu(II) to the other. Based on these data, we propose a model in which YcnJ^CopC^ and YcnI^DUF1775^ can each bind a single Cu(II) ion extracellularly and then transfer Cu(II) to the other, unmetalated protein. The YcnJ^CopC^ domain may, once Cu(II)-loaded, provide the metal to the YcnJ^CopD^ domain for uptake into the cell (**Fig. 8**).

**Fig. 8.**
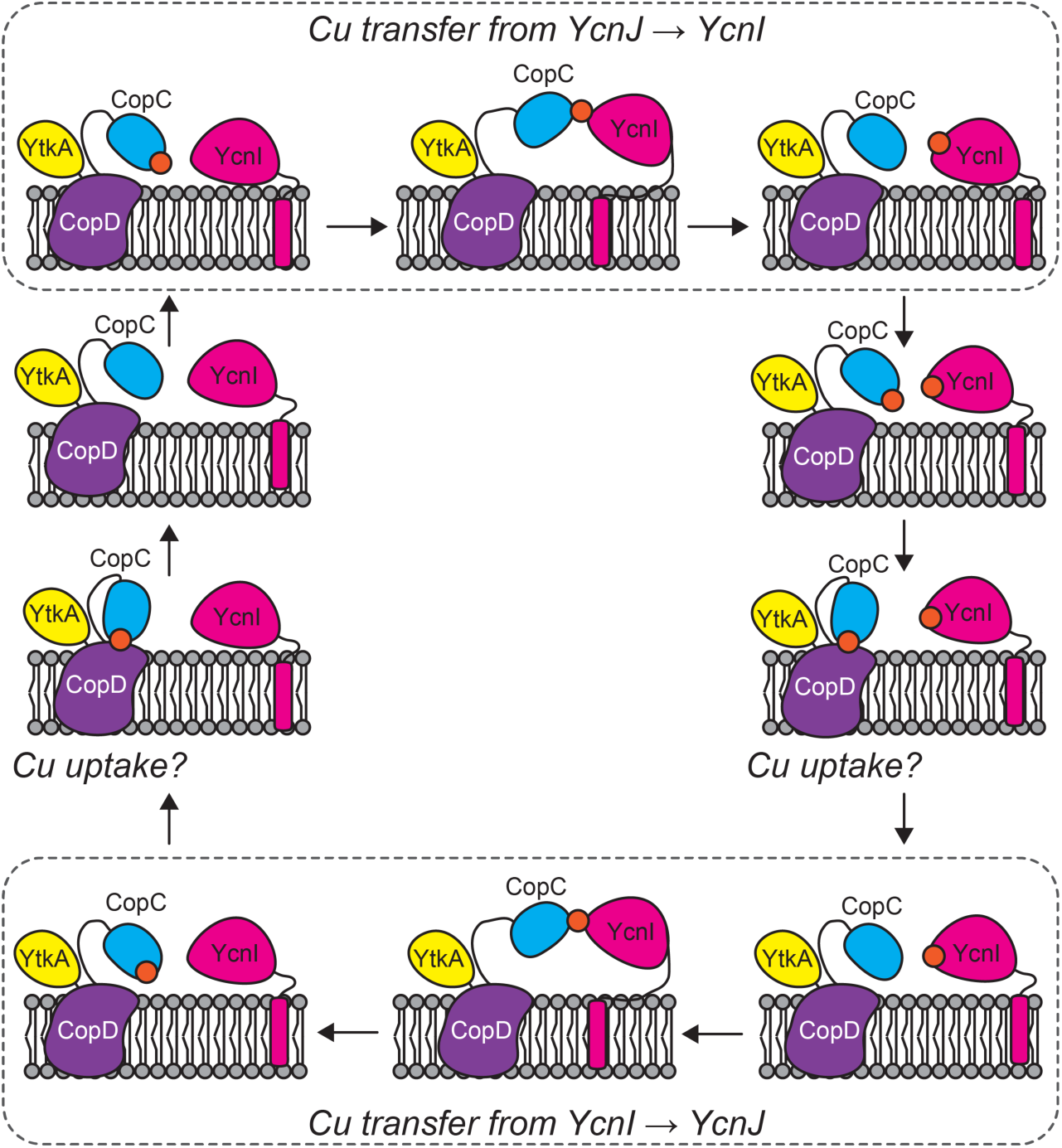
Proposed model for *in vivo* Cu(II) transfer. Cartoon depiction of proposed model for Cu transfer and subsequent uptake into the cell through CopD. YcnJ has three domains: YtkA (yellow) CopD (purple), and CopC (blue). YcnI (pink) is membrane-anchored by a C-terminal transmembrane helix. YcnJ^CopC^ and/or YcnI can bind Cu(II) (orange) extracellularly, and Cu(II) can be transferred between the CopC domains of YcnJ and YcnI (top and bottom panels). When YcnJ^CopC^ is Cu(II)-bound, it is hypothesized to present the metal to YcnJ^CopD^ for transport into the cytosol.

In our *in vitro* studies, YcnI^DUF1775^ and YcnJ^CopC^ bind Cu(II) with high affinity (3.5 fmol/L and 0.14 fmol/L, respectively) (23). While such high affinity is unusual for many biological interactions, it is quite common for Cu-binding proteins, as it reduces the amount of free copper present in cells, thereby minimizing its toxicity and allowing the organism to obtain the necessary amounts of this essential trace element when it may be scarce in the environment (1, 42, 43). Although the Cu(II) affinity of YcnJ^CopC^ is higher than that of YcnI^DUF1775^, we still observe Cu(II) transfer in both directions but with an apparent preference for transfer from YcnJ^CopC^ to YcnI^DUF1775^. Other studies of transition metal transfer between binding partners have similarly noted that the directionality of transfer does not always directly correlate to the relative affinities of both proteins (27, 34, 37, 44, 45). It is important to note that our observations of transfer *in vitro* is with the isolated YcnJ^CopC^ and YcnI^DUF1775^ domains. In the native system with intact, membrane-bound proteins that include the other domains of YcnJ, and perhaps with different relative local concentrations of the proteins, the process of Cu transfer may proceed differently from what we observe in our *in vitro* studies. Future studies of YcnJ and YcnI in their native context would therefore be informative to better understand this process.

In the case of the *ycn* system, it is possible that a transient complex is formed between YcnJ-Cu(II)-YcnI that pushes equilibrium to favor Cu(II)-YcnI over Cu(II)-YcnJ^CopC^. Perhaps this movement is electrostatically favored in this process, putting the positive charge of the Cu ion near the negatively charged π orbitals of Τrp137 and thus favoring YcnI-Cu over apoYcnI. Concurrently, in the absence of Cu, the two His ligands in YcnJ^CopC^ can engage in π-π stacking, as we observe in our crystal structure. Protonation of histidines also helps to stabilize these π-π interactions, so Cu binding may also destabilize π-π stacking (46, 47). Our data support a model in which copper can be exchanged in both directions, but that with isolated domains *in vitro*, transfer proceeds more readily from YcnJ^CopC^ to YcnI^DUF1775^.

In our *in vivo* experiments, we observe a pronounced decrease in both growth rate and yield in our Δ*ycnJ* strain under Cu limiting conditions. This is consistent with a defect in one or more Cu-dependent enzymes, which are known to include heme-copper oxidases important for energy generation during aerobic respiration. The *ycnI* deletion had little effect on growth rate in our medium conditions, but did have reduced cell yield. Strikingly, single point mutations at either of the two His that form the histidine brace motif in YcnJ^CopC^ phenocopy the *ΔycnJ* strain. Mutagenesis of YcnJ suggests that the conserved Trp137 in the monohistidine brace and membrane-localization are both important for function. Mutation of Glu50 also appeared to reduce cell yield, although we note that this residue is not as highly conserved across other DUF1775 domain sequences (22). It is curious that His27Ala had only a small growth defect under Cu-limiting conditions, but this may be due in part to the fact that YcnI plays an accessory role in Cu import and that sometimes metal-binding sites can recruit an additional ligand to compensate when another is absent. In contrast, we observe a strong phenotype when we mutate the N-terminal His in YcnJ^CopC^.

Collectively, our results suggest that the Cu(II) binding sites that we have identified *in vitro* are biologically important for Cu uptake, and that the previously functionally uncharacterized CopC and DUF1775 domains are used to facilitate and/or regulate acquisition of Cu(II). Interestingly, the copper dependent growth effects we observe in the *ycnI* gene deletion strain of *B. subtilis* hinge upon the conserved Trp residue at the copper binding site, even though we still observe copper transfer *in vitro* with this variant. Our *in vitro* experiments fail to monitor the kinetics of Cu transfer, so it is possible that the Trp residue serves to affect the rate of such a transfer event. In addition, in a biological context, both YcnI^DUF1775^ and YcnJ^CopC^ are tethered to the membrane, which likely provides additional steric constraints. It is also possible that during the transfer process, an intermediate complex forms that could destabilize the Cu-YcnJ^CopC^ interaction sufficiently to drive equilibrium towards Cu binding to YcnI.

To further consider the possibility of how the individual domains may function in the context of the full-length membrane-bound proteins, we used AlphaFold to generate predicted models for full-length YcnI, full-length YcnJ, their interactions with each other, and with Cu(II) (**Fig. S5,6**) (48–50). Interestingly, the AlphaFold predictions suggest multiple conformations for YcnJ depending on the components modeled. In the model of the apo YcnJ, the N-terminus of YcnJ^CopC^ sits in the extracellular space (**Fig. S5a, 6a**), whereas in the predicted model with Cu(II) present, the N-terminus points towards a His residue in the membrane-bound CopD domain (**Fig. S5b, 6b**). When we generate a model with YcnI, YcnJ, and a Cu(II) ion, YcnJ^CopC^ rotates further out of the membrane abutting YcnI^DUF1775^, with the Cu(II) ion predicted at their interface in a conformation that would be amenable for Cu transfer (**Fig S5c**). In the YcnI-YcnJ-Cu complex, all models have low confidence in the position of the transmembrane helix in YcnI, but consistently and with higher confidence predict the Cu ion to lie at the interface between YcnI^DUF1775^ and YcnJ^CopC^ (**Fig. S6c**). In all predictions of full-length YcnJ regardless of binding partner, there is a stretch of amino acids between the CopC and CopD domains predicted to be a disordered loop, potentially allowing YcnJ^CopC^ to swing between a conformation in which it forms a complex with YcnI and one in which it rests against the membrane.

In summary, YcnI was previously shown to bind Cu(II) (22); here we find that the extracellular CopC domain of YcnJ also binds Cu(II) in a 1:1 stoichiometry. Perhaps in the biological context the YcnJ^CopC^ domain transfers the metal ion to the CopD domain. This scenario would mean that copper reaches the importer of the *ycn* system in its oxidized Cu(II) state. Whether the metal then traverses the membrane through YcnJ being reduced *en route* through the transporter or at a later point in the process will be critical areas to pursue in the future. While these predictions could be consistent with our experimental data and our proposed model for Cu uptake (**Fig. 8**), the ability of AlphaFold to accurately predict metal binding sites is still limited. It therefore will be important for future work to experimentally determine how the full-length YcnI and YcnJ proteins interact with one another, the molecular mechanisms by which Cu is imported by *Bacillus*, and the downstream biological processes that are affected by Cu uptake.

### Experimental procedures

#### Construct design

The plasmids containing His-SUMO-YcnIΔC and His-SUMO-YcnIΔC(Trp137Phe) in pET-28a vectors were generated previously (22, 23). The His-SUMO-YcnJ^CopC^ plasmid (comprising residues 24-124 of the full-length YcnJ sequence) was synthesized commercially in the pET28a+TEV vector.

#### Protein expression and purification

The wild type and mutant versions of the soluble domain of YcnI were purified as previously described (22, 23). The His-SUMO-YcnJ^CopC^ plasmid was transformed into *Escherichia coli* (*E. coli*) BL21 (DE3) cells. An overnight culture was inoculated into Lysogeny Broth (LB) supplemented with kanamycin at 37 °C until *A*_600_ = 0.6. Protein expression was induced with 0.5 mmol/L isopropyl β-D-thiogalactopyranoside (IPTG) and grown overnight at 18 °C and 3.33 Hz (200 rpm). Cells were harvested by centrifugation at 10 000 × *g* for 30 min at 4 °C. The pellets were resuspended in pre-chilled lysis buffer (150 mmol/L NaCl, 20 mmol/L 4-(2-hydroxyethyl)piperazine-1-ethane-sulfonic acid (HEPES), 20 mmol/L imidazole, pH 7.5) supplemented with 1 mmol/L dithiothreitol (DTT), 1 mmol/L phenylmethylsulfonyl fluoride (PMSF), and 1 000 units DNase I. The resuspended pellets were sonicated on ice in 3 s / 10 s on/off cycles, for approximately 30 min and centrifuged at 15 000 × *g* for 1 h at 4 °C. The resulting clarified lysate was applied to a nickel-nitrilotriacetic acid (Ni-NTA) resin pre-equilibrated with lysis buffer (150 mmol/L NaCl, 20 mmol/L HEPES, 20 mmol/L imidazole, pH 7.5). The column was then washed with five column volumes of lysis buffer and His-SUMO-YcnJ^CopC^ was eluted with three column volumes of elution buffer (150 mmol/L NaCl, 20 mmol/L HEPES, 250 mmol/L imidazole, pH 7.5). To cleave the His-SUMO tag, the eluate was incubated with Ulp1 protease and dialyzed against size exclusion chromatography (SEC) buffer (150 mmol/L NaCl, 20 mmol/L HEPES, pH 7.5) at 4 °C overnight. The cleaved protein was further purified by applying it to Ni-NTA resin and the post-cleavage flow through and wash samples were combined and concentrated down to a volume of 1 mL to 2 mL in a 5 kDa molecular weight cutoff centrifugal concentrator at 5 000 × *g* at 4 °C. The sample was further purified by size exclusion chromatography in SEC buffer. The peak fractions containing purified protein samples were collected and concentrated. Protein concentration was measured by absorbance at 280 nm using an extinction coefficient of 11 460 (mol/L)^-1^ cm^-1^.

#### Crystallization and Structure Determination of YcnJ^CopC^

Initial crystallization screens of YcnJ^CopC^ were carried out using the sitting drop method at room temperature in 96-well crystallization plates. An initial hit was obtained in 0.2 mol/L ammonium sulfate, PEG 8 000 at a mass fraction of 30 %, which was further manually refined. Crystals were obtained in 0.2 mol/L lithium sulfate, PEG 8 000 at a mass fraction of 39 % using the cross-seeding technique, with seeds obtained from crystals from YcnI Trp137Phe (PDB ID: 8UM6). Data were collected at the 21-ID-D beamline at the Advanced Photon Source and processed to 0.160 nm (1.60 Å) resolution using DIALS User Interface (DUI) (51) in space group *P*4_3_2_1_2. The structure was solved by molecular replacement using the open-source software phenix.phaser (52) from the Phenix package (53) using a model generated using the AlphaFold2-based platform ColabFold (54). The structure was further improved by iterative rounds of model building and refinement in phenix.refine and Coot (55). The final model consists of 97 residues with *R*_work_/*R*_free_ = 21.07 % / 23.27 % (**Table S1**). Software used in this project was curated by SBGrid (56).

#### Determination of Cu content by ICP-MS

All Cu(II)-binding experiments were conducted aerobically. Samples were prepared by adding 1 or 2 molar equivalents of CuSO_4_ solution to 60 μmol/L protein solution either by slow addition by pipetting or by using a syringe pump at a rate of 0.3 μL/min. The samples were incubated on ice for 1 h prior to desalting using a ZebaSpin desalting column pre-equilibrated in SEC buffer to remove free Cu ions. Protein samples were digested in water containing a nitric acid volume fraction of 1 % to a final protein concentration of 0.1 μmol/L. Total copper was quantified using quadrupole inductively coupled plasma mass spectrometry with pneumatic nebulization and spraychamber cooled to 2 °C. The instrument was optimized and calibrated using ICP Mix Standard 5 (Inorganic Ventures, see Disclaimer) to measure copper concentrations over a working range of 1.6 nmol/L to 400 nmol/L.

#### Continuous-wave (CW) EPR spectroscopy

Approximately 300 µL of each sample (400 µmol/L Cu-loaded YcnJ^CopC^, 400 µmol/L apo YcnJ^CopC^, and buffer) was loaded into quartz EPR tubes with an outer diameter of 4 mm for low T (80 K) CW measurements. Samples were capped, frozen to −80 °C, and transferred to liquid N_2_ prior to placement in the cryostat. Spectra were collected on a commercial EPR spectrometer operating at 9 GHz (X-band). The temperature was maintained at 80 K using liquid nitrogen, a commercially available cryostat, and a temperature control system designed for low temperature CW EPR spectroscopy. All spectra were collected with a modulation amplitude of 0.2 mT and microwave powers of 2.4 mW or 0.4 mW as indicated in the figure legends. Spectra with a scan width of 200 mT were collected with 2048 points, conversion time of 58.59 ms, sweep time of 2 min, and either 4 scans or 16 scans as indicated in the figure legends. Spectra with a scan width of 17 mT were collected with 512 points, conversion time 117.19 ms, sweep time of 1 min, and 4 scans. The temperature was (80 +/− 0.8) K for all spectra with the uncertainty on the temperature representing the maximum deviation of the measurement temperature from the setpoint (80.0 K) during the scan.

#### Cu(II) affinity measurements by Isothermal Titration Calorimetry (ITC)

All ITC experiments were conducted using a low volume isothermal titration calorimeter at 25 °C. Glycine was used as a competitor for copper to accurately measure the high affinity protein-copper interaction following a previously described protocol (57), and its usefulness as a weaker chelator in Cu(II) binding measurements by ITC has been demonstrated (57–59). Protein samples were desalted into 150 mmol/L NaCl, 20 mmol/L HEPES, 30 mmol/L glycine, pH 7.5 using a ZebaSpin desalting column. CuSO_4_ was prepared in the same buffer at 500 μmol/L. The sample cell was loaded with 50 μmol/L apo YcnJ^CopC^, and or 500 μmol/L CuSO_4_ was loaded in the injection syringe. An initial injection of 0.4 μL CuSO_4_ was followed by injections each with a volume of 2.0 μL and constant stirring at 12.5 Hz (750 rpm). The total number of injections was 17 and the spacing between injections was 150 s. Experiments were performed in duplicate, and a blank run injecting CuSO_4_ into buffer was subtracted from data. The resulting data were analyzed using commercial graphing software and fit to a one-site binding model. Conditional *K*_D_ values considering the presence of glycine in the reaction were derived as described in (58).

#### Bacillus strain construction and growth conditions

Mutant alleles of the *ycnI* and *ycnJ* genes were generated using the simplified Clustered Regularly Interspaced Short Palindromic Repeats (CRISPR)-cas9 method for *B. subtilis* as described previously (60). The repair fragments containing the desired mutation were amplified by long-flanking homology (LFH) polymerase chain reaction (PCR), with appropriate upstream and downstream homology using primers containing the SfiI restriction sites. These fragments were then joined by a stitching PCR using commercially available, high-fidelity DNA polymerase. The purified stitched PCRs were digested with SfiI (New England Biolabs, NEB) at 50 °C for 2 h along with the pAJS23 plasmid which encodes the guide RNA that recognizes the *erm* cassette (60). The digested, column purified (OMEGA) repair templates were cloned into the pAJS23 plasmid using T4 DNA ligase (NEB). These ligated products were then transformed into *E. coli* DH5α and selected on LB agar plates supplemented with kanamycin (30 μg/mL) for propagation and sequence confirmation. Correct clones were identified using Sanger sequencing (BRC at Cornell). Further, plasmid was moved into *E. coli* TG1 background. Plasmid DNA was transformed into *ycnJ*::*erm* or *ycnI*::*erm* (NRB, Japan) recipient strain of *Bacillus subtilis.* Briefly, the *erm-*knockout strains were aerobically grown in 5 mL of modified competence (MC) media at 37 °C to optical density (OD 600 nm) of 0.8. Then, cultures were shifted to 30 °C and 1 mg of plasmid DNA was added. Cultures were incubated for 2 h and 100 µL of transformation reactions were plated onto LB agar containing kanamycin (15 μg/mL) plus 0.011 mol/L (0.2 % w/v) mannose. These plates were incubated at 30 °C for 48 h. To cure the plasmid, transformants were repeatedly passaged by patching onto LB agar (without any antibiotics) for 3 days at 45 °C. Finally, patch plates were made on LB, LB plus erythromycin, LB plus kanamycin; of these plates only the erythromycin and kanamycin sensitive clones were selected indicating successful replacement of the *erm* cassette. The resulting strains were confirmed by Sanger sequencing to verify the presence of the desired mutations. The strains are listed in **Table S2**.

#### Growth conditions

Bacterial cultures were streaked from frozen glycerol stocks on LB agar plates overnight at 37 °C. Cells were transferred to 5 mL of LB broth and were aerobically grown at 37 °C until *A*_600_ reached approximately 0.4. From these mid-log cultures, 1 mL was transferred to a microcentrifuge tube and washed with a defined, minimal media malate (MM-malate). In a 100 well honeycomb microplate, 200 μL of MM-malate was dispensed and to this 2 μL of bacterial inoculum was added. In Cu-depleted conditions, 10 mmol/L of triethylenetetramine (TETA) was added to the MM-malate medium. Growth was monitored periodically for 48 h with shaking at 37 °C using a photometric microplate absorbance reader and growth curve analyzer. The growth curve shows 3 biological replicates with 2 technical replicates.

#### Cu transfer experiments

All Cu(II)-binding experiments were conducted aerobically. 500 µL CuSO_4_ in ion exchange (IEX) buffer A (20 mmol/L HEPES, pH 7.5) was slowly pipetted over an approximately 30 s to 1 min timeframe into 500 µL concentrated protein samples in a 2:1 molar ratio. The mix was incubated on ice for 1h and then diluted to 2 mL in pre-chilled IEX buffer A. Solution was centrifuged at 10 000 x *g* for 10 min at 4 °C before being injected on a HiTrap Q XL 1 mL column. Cu-loaded protein was separated from apo protein and free Cu ions by anion exchange chromatography and copper content of fractions were measured by the BCA assay.

A 1 mL solution containing 100 µmol/L Cu-loaded protein and 100 µmol/L apo protein was incubated on ice for 30 min. For control experiments, a 1 mL solution containing 100 µmol/L Cu-loaded protein was incubated in SEC buffer. Solutions were diluted to 2 mL in pre-chilled IEX buffer A. Solution was centrifuged at 10 000 x g for 10 min at 4 °C before being injected on a HiTrap Q XL 1 mL column.

Proteins were separated by anion exchange chromatography and copper content of fractions were measured by the BCA assay, as follows. A 3 mmol/L stock of bicinchoninic acid disodium salt (BCA) was prepared in SEC buffer and a 0.1 mol/L ascorbate solution was prepared in deionized H_2_O. All stocks were freshly prepared before each experiment. Protein samples at 100 µmol/L were treated with 1.5 mmol/L ascorbic acid to reduce Cu(II) to Cu(I) and incubated for approximately 10 min. The BCA solution was added to each sample in a 1:3 dilution and spectra from 400 nm to 700 nm were measured on a double-beam, double-monochromator UV/visible/near infrared spectrophotometer. The concentration of BCA-Cu complex was calculated based on absorbance at 562 nm using an extinction coefficient of 7 900 (mol/L)^-1^ cm^-1^ (61). Protein concentrations of the resulting fractions were measured by absorbance at 280 nm to calculate Cu:protein stoichiometries. Each experiment was conducted 3 to 4 times.

For the dialysis experiments, Cu-loaded proteins or as-purified proteins were prepared as above. Cu-loaded protein (250 µL) was loaded into a 3.5kDa MWCO dialysis cassette. The as-purified protein (250 µL) was loaded into a second 3.5 kDa MWCO dialysis cassette. The two cassettes were placed in a beaker containing 150-200 mL SEC buffer and was dialyzed overnight at 4 ℃. The following morning, the proteins were removed from the cassettes and copper content was determined by the BCA assay. Each experiment was conducted 3 times.

## Data availability

The final coordinates and structure factors for the YcnJ^CopC^ structure have been deposited in the PDB as 9C14 and the dataset has been deposited in the SBGrid database as 10.15785/SBGRID/1108. All other data are available upon request.

## Supporting Information

This article contains supporting information.

## Acknowledgements

We thank Prof. Ronen Marmorstein and Kollin Schultz (University of Pennsylvania) for use of the ITC instrument, Dr. Wendy Breyer (Lehigh University) for assistance with instrumentation, and members of the Fisher laboratory for helpful discussions.

## Disclaimer

Certain commercial firms and trade names are identified to specify the usage procedures adequately for reproducibility. Such identification is not intended to imply recommendation or endorsement by the National Institute of Standards and Technology, nor is it intended to imply that related products are necessarily the best available for the purpose.

