## Supplemental Information for "Copper acquisition in *Bacillus subtilis* involves Cu(II) exchange between YcnI and YcnJ"

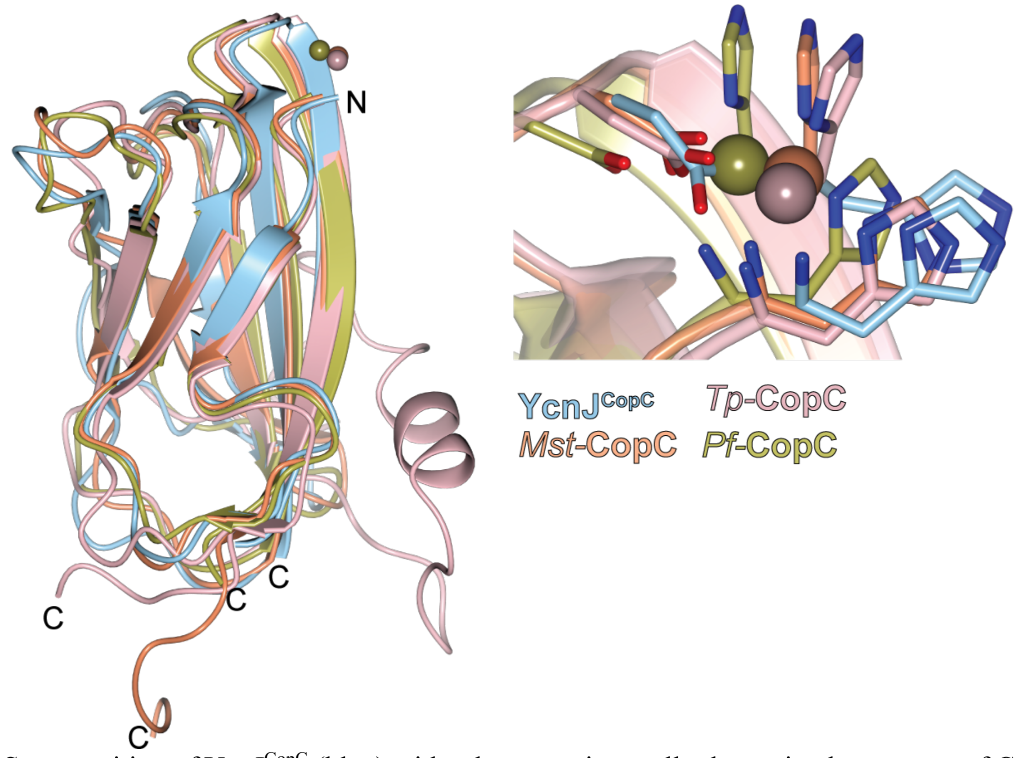


**Fig. S1.** Superposition of YcnJ$^{CopC}$ (blue) with other experimentally determined structures of Cu(II)-bound CopC proteins (PDB IDs 5ICU – from *Methylosinus trichosproium OB3b* [13], 6NFQ – from *Pseudomonas fluorescens* [17], 8YTR – from *Thioalkalivibrio paradoxus Arh1*) and their Cu(II)-binding sites as an inset.

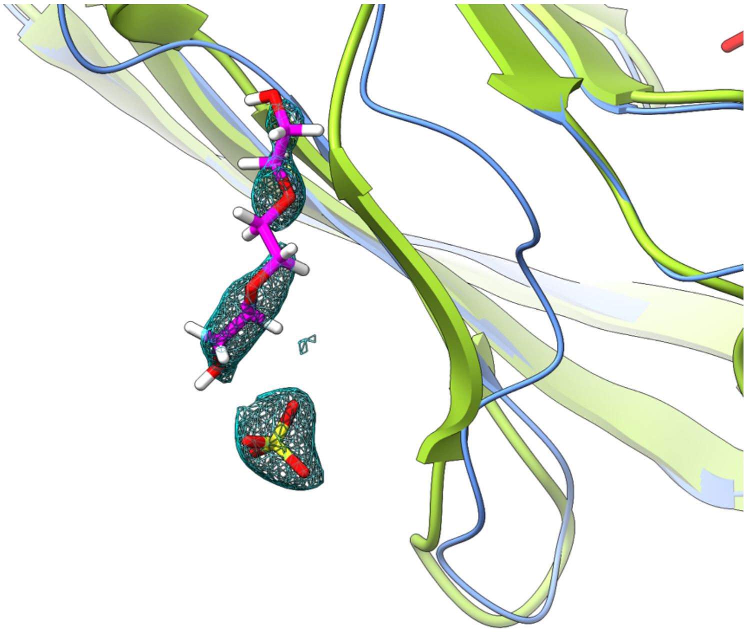

**Figure S2. Strand displacement is a crystallographic artifact.** A β-strand present in other CopC proteins, such as that from *M. trichosporium OB3b* (PDB ID 5ICU, yellow green) is displaced by the presence of triethylene glycol and a sulfate ion (magenta and yellow, respectively) in the structure of $YcnJ^{CopC}$ (cornflower blue). $2F_o$-$F_c$ maps of triethylene glycol and $SO_4$ are shown (contour level: 0.30).

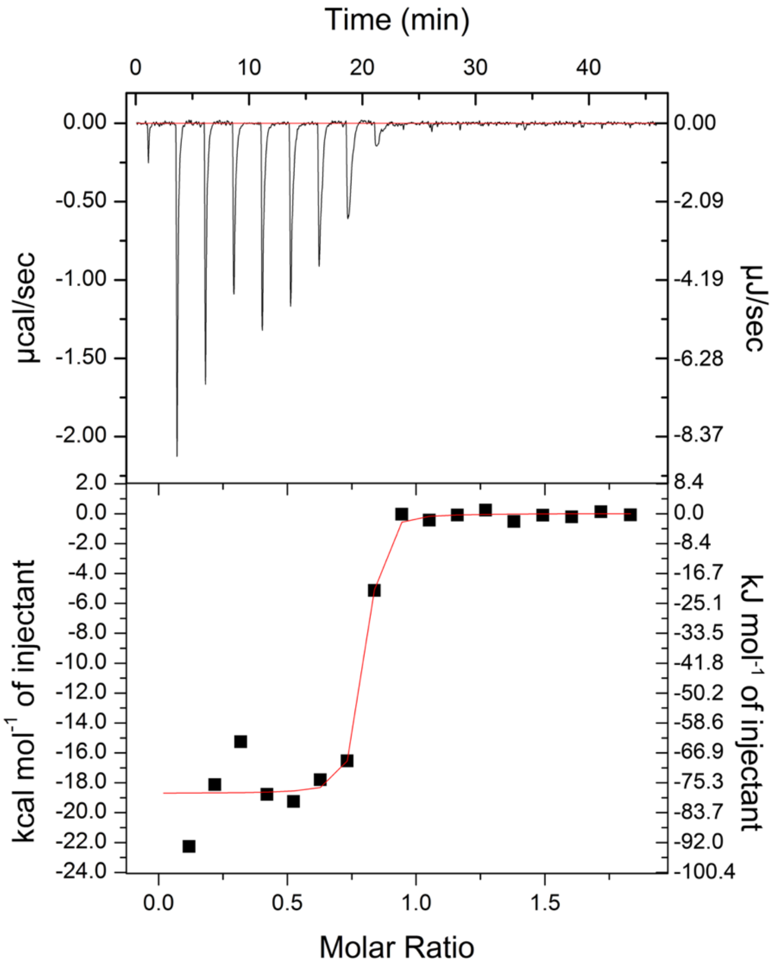


**Figure S3. Replicate experiment of isothermal titration calorimetry of Cu(II) into YcnJ$^{CopC}$.** Measurements were performed in the presence of 30 mmol/L glycine as a weak competitor. Conditional $K_D$ = 1.32 x $10^{-16}$ mol/L; $n_{ITC}$ = 0.752 ± 0.0121; ΔH = -78 212 ± 2 495 J/mol; ΔS = −120 J/mol/K. Reported error represents error on the curve fit.

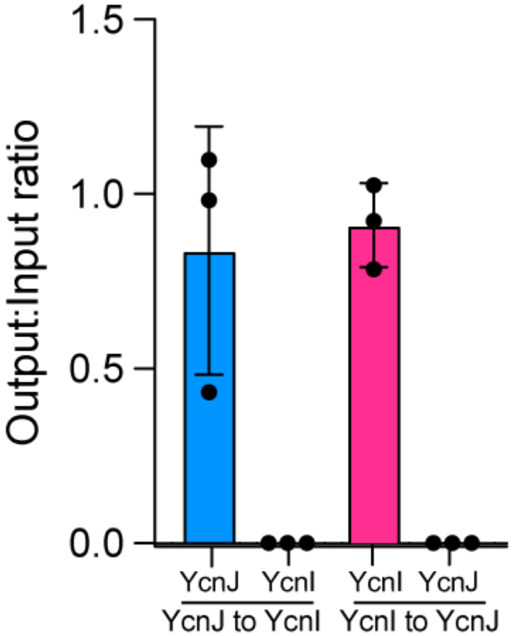


**Fig. S4. Cu cannot passively transfer between YcnJ and YcnI.** Average ratios of Cu(II) concentrations before and after dialysis of Cu(II)-YcnJ$^{CopC}$ with as-purified YcnI$^{DUF1775}$ (blue) and Cu(II)-YcnI$^{DUF1775}$ with as-purified YcnJ$^{CopC}$ (magenta). For as-purified proteins, only the results after dialysis are shown. The experiment was repeated 3 times. Individual results are shown as black circle and error bars represent calculated SD.

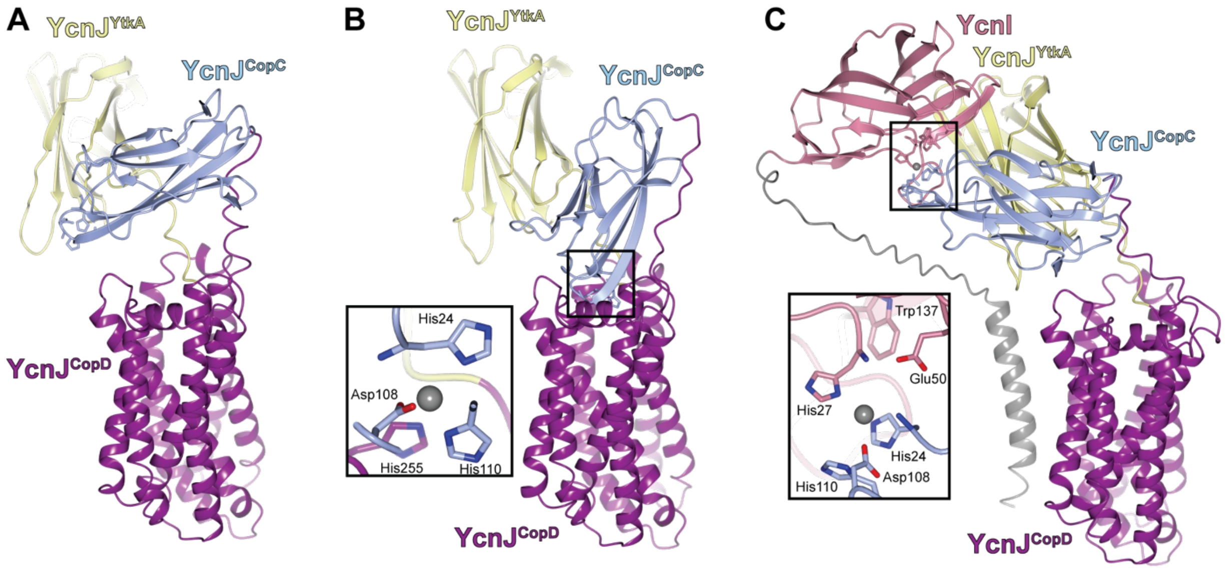


**Figure S5. Structural predictions of full-length YcnJ.** AlphaFold3 predictions for (a) full-length apo YcnJ, (b) full-length YcnJ with 1 Cu(II) ion, and (c) full-length YcnJ in complex with full-length YcnI and 1 Cu(II) ion. Proteins are colored by domain, and insets represent predicted locations for the Cu(II) ion.

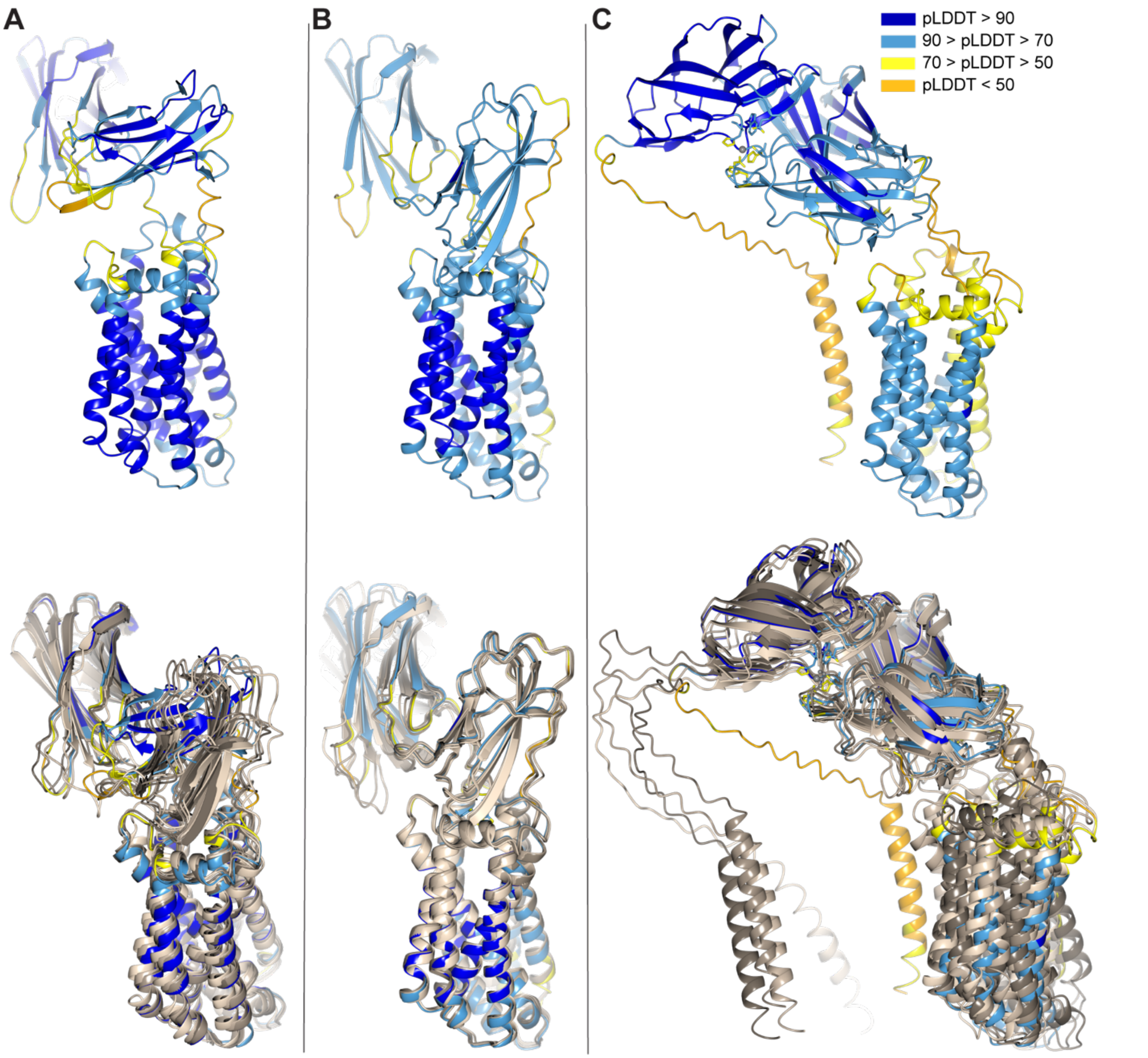


**Figure S6. AlphaFold3 predictions.** The same AlphaFold3 models from Fig. S4 colored by pLDDT values (top) and superposition of the five best models output (bottom). (A) full-length apo YcnJ, (B) full-length YcnJ with 1 Cu(II) ion, and (C) full-length YcnJ in complex with full-length YcnI and 1 Cu(II) ion.

**Table S1. Data collection and refinement statistics**

| Crystal | $YcnJ^{CopC}$ |
|---|---|
| PDB accession code | 9C14 |
| Data collection | |
| Wavelength (Å) | 1.378 |
| Space group | $P4_32_12$ |
| Cell dimensions | |
| a, b, c (Å) | 36.0, 36.0, 165.8 |
| α, β, γ (°) | 90.0, 90.0, 90.0 |
| Resolution (Å)* | 41.44 to 1.60 (1.63 to 1.60) |
| $R_{meas}$* | 0.876 (40.355) |
| $CC_{½}$* | 0.997 (0.516) |
| I / σI* | 9.6 (1.00) |
| $R_{merge}$* | 0.858 (-) |
| $R_{pim}$* | 0.177 (8.103) |
| Completeness (%)* | 100.0 (100.0) |
| Redundancy* | 24.1 (24.6) |
| Refinement | |
| Resolution (Å)* | 41.44 to 1.60 (1.65 to 1.60) |
| No. of reflections* | 15 313 (1 209) |
| $R_{work}$ / $R_{free}$ (%)* | 21.1 / 23.3 (47.5 / 48.6) |
| Residue range built | 24 to 120 |
| No. of atoms | |
| Protein | 789 |
| Ligand/ion | 10 $SO_4$, 24 PGE |
| Water | 62 |
| Model Quality | |
| B-factors ($Å^2$) | |
| Overall | 24.00 |
| Protein | 35.55 |
| Ligand/ion | 46, 48, 30 |
| Water | |
| RMSD, bond lengths (Å) | |
| RMSD, bond angles (°) | |
| Ramachandran favored/allowed/outliers (%) | 95 / 2 / 0 |

*Parentheses indicate highest resolution shell.
Note that 1 nm = 10 Å

**Table S2. Strains used in this study**

| Strain | Genotype | Construction | Reference |
|---|---|---|---|
| ***B. subtilis*** | | | |
| CU1065 | *WT* | Lab strain | Lab stock |
| HB30921 | *ΔycnJ::erm* | BGSC gDNA-->CU1065 | This work |
| HB30922 | *ΔycnI::erm* | BGSC gDNA-->CU1065 | This work |
| HB30927 | *ΔycnJ* | pDR244-->HB30921 | This work |
| HB30930 | *ΔycnI* | pDR244-->HB30922 | This work |
| HB30956 | *ycnI (His27Ala)* | CRISPR (pAJS23+repair template)--> HB30922 | This work |
| HB30958 | *ycnI (Glu50Ala)* | CRISPR (pAJS23+repair template)--> HB30922 | This work |
| HB30959 | *ycnI (Trp137Phe)* | CRISPR (pAJS23+repair template)--> HB30922 | This work |
| HB30927 | *ycnI (TruncAsp170)* | CRISPR (pAJS23+repair template)--> HB30922 | This work |
| HB30960 | *ycnJ (His24Ala)* | CRISPR (pAJS23+repair template)--> HB30921 | This work |
| HB30977 | *ycnJ (His110Ala)* | CRISPR (pAJS23+repair template)--> HB30921 | This work |

**Table S3. Conditional Dissociation Constants ($K_D$) of binding of CopC proteins to Cu(II).**

| Protein | $K_D$ (M) | n | Reference |
|---|---|---|---|
| YcnJ$^{CopC}$ (1) | 1.43 x $10^{-16}$ | 0.682 | This study |
| YcnJ$^{CopC}$ (2) | 1.32 x $10^{-16}$ | 0.752 | This study |
| YcnI$^{WT}$ | 3.51 x $10^{-15}$ | 1.07 | de Oliveira Silva et al. (23) |
| YcnI$^{W137F}$ | 2.02 x $10^{-14}$ | 0.35 | de Oliveira Silva et al. (23) |
| YobA | 3 x $10^{-9}$ | 1.09 | Hadley et al. (15) |
| *Pf*CopC* | $10^{-16}$ | - | Wijekoon et al (20) |
| *Ps*CopC* | $10^{-14}$ | - | Wijekoon et al. (20)<br>Zhang et al. (26) |

*Constants measured via ligand competition using fluorescent probes.
